# Incidence of an intracellular multiplication niche amongst *Acinetobacter baumannii* clinical isolates

**DOI:** 10.1101/2021.04.15.439986

**Authors:** Tristan Rubio, Stéphanie Gagné, Charline Debruyne, Chloé Dias, Caroline Cluzel, Doriane Mongellaz, Patricia Rousselle, Stephan Göttig, Harald Seifert, Paul G. Higgins, Suzana P. Salcedo

## Abstract

The spread of antibiotic resistant *Acinetobacter baumannii* poses a significant threat to public health worldwide. This nosocomial bacterial pathogen can be associated with life-threatening infections, particularly in intensive care units. *A. baumannii* is mainly described as an extracellular pathogen with restricted survival within cells. This study shows that a subset of *A. baumannii* clinical isolates extensively multiply within non-phagocytic immortalized and primary cells, without the induction of apoptosis, and with bacterial clusters visible up to 48 hours after infection. This phenotype was observed for the *A. baumannii* C4 strain associated with high mortality in a hospital outbreak, and the *A. baumannii* ABC141 strain which wasn’t isolated from an infection site but was found to be hyperinvasive. Intracellular multiplication of these *A. baumannii* strains occurred within spacious single membrane-bound vacuoles, labeled with the lysosomal associate membrane protein (LAMP1). However, these compartments excluded lysotracker, an indicator of acidic pH, suggesting that *A. baumannii* can divert its trafficking away from the lysosomal degradative pathway. These compartments were also devoid of autophagy features. A high-content microscopy screen of 43 additional *A. baumannii* clinical strains highlighted various phenotypes: (1) the majority of strains remained extracellular, (2) a significant proportion was capable of invasion and limited persistence, and (3) two strains efficiently multiplied within LAMP1-positive vacuoles, one of which was also hyperinvasive. These data identify an intracellular niche for specific *A. baumannii* clinical strains that enables extensive multiplication in an environment protected from host immune responses and out of reach from many antibiotics.

**Importance:** Multidrug resistant *Acinetobacter baumannii* strains are associated with significant morbidity and mortality in hospitals world-wide. Understanding their pathogenicity is critical for improving therapeutics. Although *A. baumannii* can steadily adhere to surfaces and host cells, most bacteria remain extracellular. Recent studies have shown that a small proportion of bacteria can invade cells but present limited survival. We have found that some *A. baumannii* clinical isolates can establish a specialized intracellular niche that sustains extensive intracellular multiplication for a prolonged time without induction of cell death. We propose that this intracellular compartment allows *A. baumannii* to escape the cell’s normal degradative pathway, protecting bacteria from host immune responses and potentially hindering antibiotic accessibility. This may contribute to *A. baumannii* persistence, relapsing infections and enhanced mortality in susceptible patients. A high-content microscopy-based screen confirmed this pathogenicity trait is present in other clinical isolates. There is an urgent need for new antibiotics or alternative antimicrobial approaches, particularly to combat carbapenem-resistant *A. baumannii*. The discovery of an intracellular niche for this pathogen as well as hyperinvasive isolates may help guide the development of antimicrobial therapies and diagnostics in the future.

## Introduction

*Acinetobacter baumannii* is a nosocomial pathogen posing a growing global health threat due to its remarkable ability to persist in the environment and acquire extensive multi-drug resistance. In some countries, carbapenem resistance rates have surpassed 80% (1) ranking this pathogen as a top priority for developing new antibiotics by the World Health Organization (2). Carbapenem resistance is associated mostly with eight international clonal (IC) lineages (3). Although community-acquired cases have been described, *A. baumannii* mainly impacts patients with severe underlying disease such as those in intensive care units. One of the most frequent clinical manifestations of *A. baumannii* infection is ventilator-associated pneumonia (VAP), often associated with a poor prognosis. Of increasing concern is the recent appearance of hypervirulent strains that present concurrently extensive antibiotic resistance and have been implicated in hospital and animal infection outbreaks of which some were fatal (4–6). Despite its growing importance, the mechanisms underlying *A. baumannii* virulence remain poorly characterized. Its ability to adhere to abiotic surfaces and form biofilms enables colonization of medical equipment and surfaces (7). Adherence to human cells and the interplay with innate immune cells have also proven critical to *A. baumannii* virulence (8, 9). *A. baumannii* is primarily considered as an extracellular pathogen. In some studies, clinical strains were described as non-invasive in human lung epithelial cell lines (10). *A. baumannii* laboratory and clinical strains were also shown to be rapidly phagocytosed and killed by cultured macrophages and neutrophils (11, 12). However, previous studies have highlighted the ability of different A. baumannii strains to be internalised or to actively invade host cells (13–20). Intracellular survival of *A. baumannii* in cultured cells has been reported when critical anti-bacterial host response pathways were inhibited, such as Nod1/Nod2, nitric oxide or autophagy (12, 14, 15). A few recent studies have suggested that some strains of *A. baumannii* can invade and transiently survive within epithelial human cells and macrophages (16–18, 20, 21). Although the *A. baumannii* strain ATCC 19606 is killed by macrophages, it was shown to enter epithelial cells by a zipper-like mechanism associated with actin microfilaments and microtubules (16). Similarly, the *A. baumannii* strain ATCC 17978 can survive within human epithelial lung cells resulting in activation of lysosomal biogenesis and autophagy (18). More recently, the strain *A. baumannii* AB5075-UW was also shown to invade non-phagocytic cells by binding carcinoembryonic antigen-related cell adhesion (CECAM) molecules (17). Once intracellular, AB5075-UW survive within a vacuole associated with early and late endosomal GTPases Rab5 and Rab7 as well as the autophagy protein light chain 3 (LC3). Nevertheless, bacteria are progressively killed by vacuolar acidification (17). In this work, we highlight several *A. baumannii* clinical isolates that multiply intracellularly within large late-endosomal-derived vacuoles without autophagy features. We found that this intracellular multiplication is not associated with cytotoxicity. High-content screening of 42 clinical isolates suggests that a significant proportion of isolates are capable of invasion and intracellular survival, with a minor subset able to establish intracellular replication niches. Notably, a few strains were hyper-invasive and hyper-replicative. Taken together, these results shed some light on a potentially clinically relevant intracellular niche for some *A. baumannii* isolates which could impact patient management in a hospital setting, provide a target for new therapeutic approaches or constitute a biomarker for virulent strains.

## Results

### The hypervirulent C4 clinical strain is able to invade human A549 lung epithelial cells

Given the increasing reports on hypervirulent strains of *A. baumannii* in hospitals, we set out to investigate if this enhanced virulence could be attributed to particular interactions with host cells. We initially focused on the *A. baumannii* strain C4 belonging to the international clonal lineage 4 (IC4), isolated from a wound swab from a patient hospitalized in Germany in in 2010. This isolate was transmitted between 4 patients who subsequently died, and we hypothesized that this strain was hypervirulent. We first assessed the virulence of this strain *in vitro* using the well-established *Galleria mellonella* infection model. The experimental infection revealed that C4 is significantly more virulent than the well-characterized *A. baumannii* strain ATCC 17978, used as a control in this study (Fig. 1A). To investigate the mechanisms underlying the virulence of the C4 strain, we infected human lung epithelial A549 cells and quantified the levels of adhesion at 1 h post-infection (p.i.). Analysis of the percentage of bacterial adhesion relative to the inocula was equivalent for both revealed that the C4 strain does not have enhanced adhesion capacity (Fig. 1B). We next measured the levels of LDH released from cells infected for 6h with *A. baumannii* C4 in comparison with ATCC 17978, the environmental *A. baumannii* strain DSM 30011 and the cytotoxic *Pseudomonas aeruginosa* strain PA14. Cells were either infected for 1h and then incubated with antibiotics to eliminate extracellular bacteria or washed to remove non-adherent bacteria without the use of antibiotics. No significant cytotoxicity was observed in cells infected with any of the *A. baumannii* strains tested, in contrast to P. aeruginosa (Fig. 1C). We next assessed the production of the proinflammatory interleukin IL6 by A549 cells infected with *A. baumannii* C4, ATCC 17978, and DSM30011 at 3, 8 and 24 h p.i. As previously reported, ATCC 17978 induced IL6 secretion with a peak at 24 h p.i. (Fig. 1D). Similar levels of IL6 were detected in cells infected with the environmental *A. baumannii* isolate DSM 30011. Interestingly, the C4 strain seemed to result in a lower increase of IL6 production between 8 and 24 h p.i., suggesting that the inflammatory response to C4 may be less pronounced than the response elicited by ATCC 17978 and DSM 30011 (Fig. 1D). Finally, we tested whether C4 can invade human non-phagocytic cells. We observed a clear condensation of actin around the bacteria upon entry into the cell of the C4 strain, suggesting that actin cytoskeleton re-arrangements are involved in the invasion process (Fig. 1E) as previously described for the strain ATCC 19606 (16). At 24 h p.i., confocal microscopy revealed intracellular clusters of bacteria suggestive of intracellular replication (Fig. 1F). Z-stack analysis of infected cells labelled with tubulin confirmed the intracellular nature of these bacterial clusters (Fig. 1F). In summary, the *A. baumannii* C4 strain can invade host cells and form intracellular bacterial clusters without causing cell lysis.

**Fig. 1.**
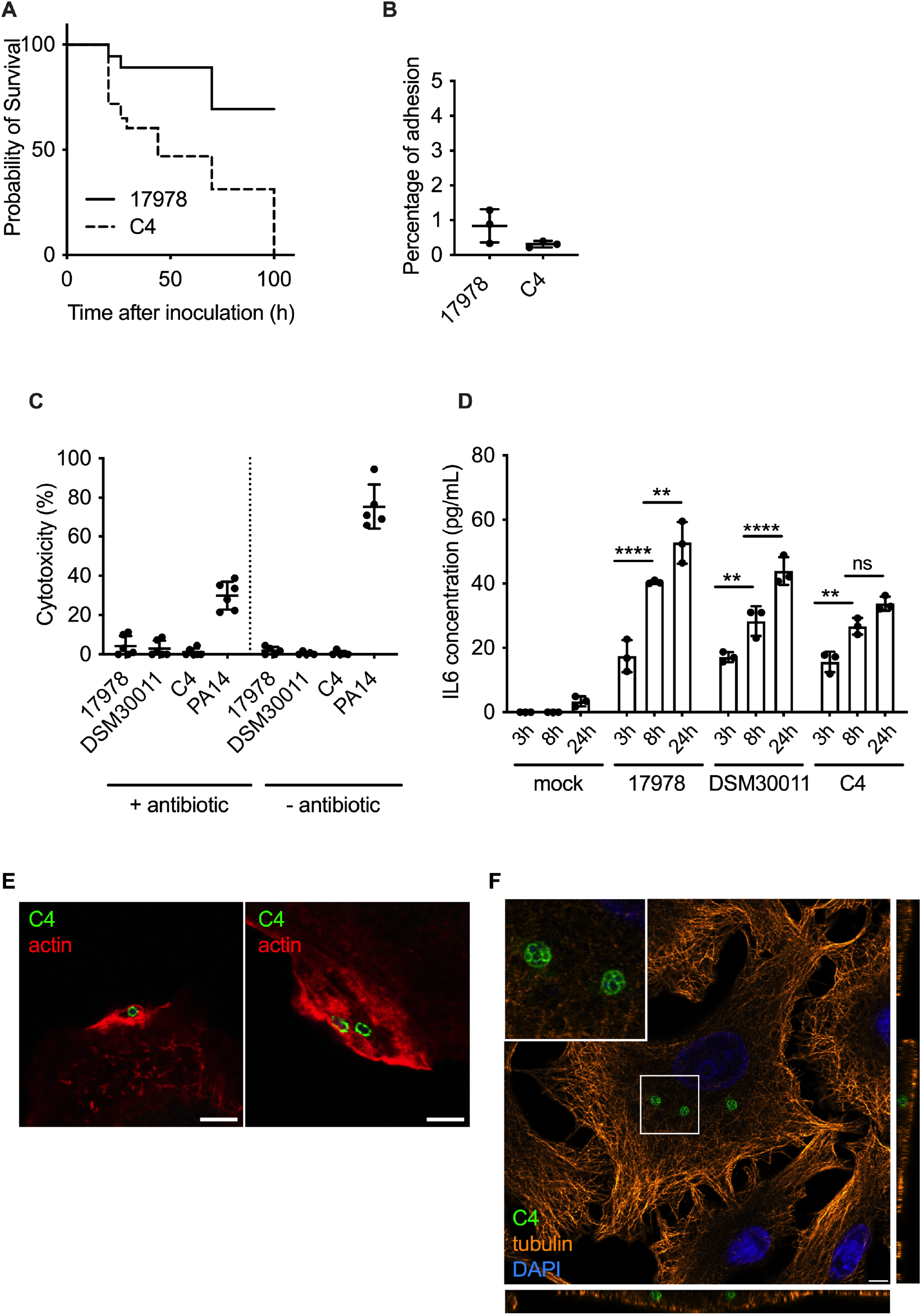
Characterization of the hypervirulent *A. baumannii* C4 clinical strain. **(A)** Kaplan–Meier survival curves were generated from *G. mellonella* injected with the *A. baumannii* strains C4 or ATCC 17978 (1×10^6^ CFU per insect). Mortalities were counted regularly over 100 hours. *A. baumannii* C4 is significantly more virulent than ATCC 17978. **** *p*<0.0001 (log-rank test). Data are representative of 3 independent experiments. **(B)** Percentage of C4 and 17978 adhesion to A549 cells (MOI 100:1). Data correspond to mean *±* SD, from 3 independent experiments. Their percentage of adhesion are not significantly different. **(C)** Cytotoxicity of *A. baumannii* ATCC 17978, DSM 30011, C4 and *P. aeruginosa* PA14 were monitored using LDH assay in A549 cells in the presence or absence of antibiotic. No cytotoxicity was observed for any *A. baumannii* strains. Data correspond to mean *±* SD, from 5 independent experiments. **(D)** Quantification of IL6 concentration produced by A549 cells infected by *A. baumannii* ATCC 17978, DSM 30011 and C4 at 3 h, 8 h and 24 h p.i. Statistical comparison was done with two-way ANOVA with a Holm-Sidak’s correction for multiple comparisons. Between *A. baumannii* ATCC 17978 3h and 8h (****) *p*<0.0001, between DSM 30011 3h and 8h (**) p<0.01, between C4 3h and 8h (**) *p*<0.01, between ATCC 17978 8h and 24h (**) *p*<0.01 and between DSM 30011 8h and 24h (****) *p*<0.0001. Not all comparisons are shown. Data correspond to mean *±* SD, from 3 independent experiments. **(E)** A549 cells were infected with C4, immunolabelled 1 h p.i. and analyzed using confocal immunofluorescence microscopy. *A. baumannii* (green) was labelled with specific antibodies and phalloidin was used to visualize actin (red). Scale bars correspond to 5 μm. **(F)** A549 cells were infected with C4 and labelled for tubulin (orange), *A. baumannii* (green) and the nucleus with DAPI (blue). The orthogonal view of the z-stack show that C4 is able to enter into the cell and form intracellular bacterial clusters.

### Clinical *A. baumannii* strains C4 and ABC141 multiply in human cells

To determine if the presence of intracellular bacterial clusters at 24 h p.i. resulted from intracellular replication by the *A. baumannii* C4 strain, we quantified by microscopy the number of bacteria per cell at 1 and 24 h p.i. for the C4 and ATCC 17978 strains. We initially tested two bacterial growth conditions, exponential and stationary. Equivalent results were obtained for both bacterial growth phases, so only data referring to exponential growth is shown, and we selected this condition for all subsequent experiments. At 1h, we observed that infected cells contained one or two bacteria per cell for either the C4 and ATCC 17978 strains (Fig. 2A) with equivalent percentages of infected cells observed (Fig. 2B). At 24 h p.i. we could detect the appearance of bacterial clusters with multiple bacteria (up to 20 bacteria per cluster) for the *A. baumannii* C4 strain in contrast to the ATCC 17978 which remained with only a few bacteria per cell (Fig. 2A and C). Together these results indicate that the virulent *A. baumannii* C4 strain is capable of intracellular multiplication.

**Fig. 2.**
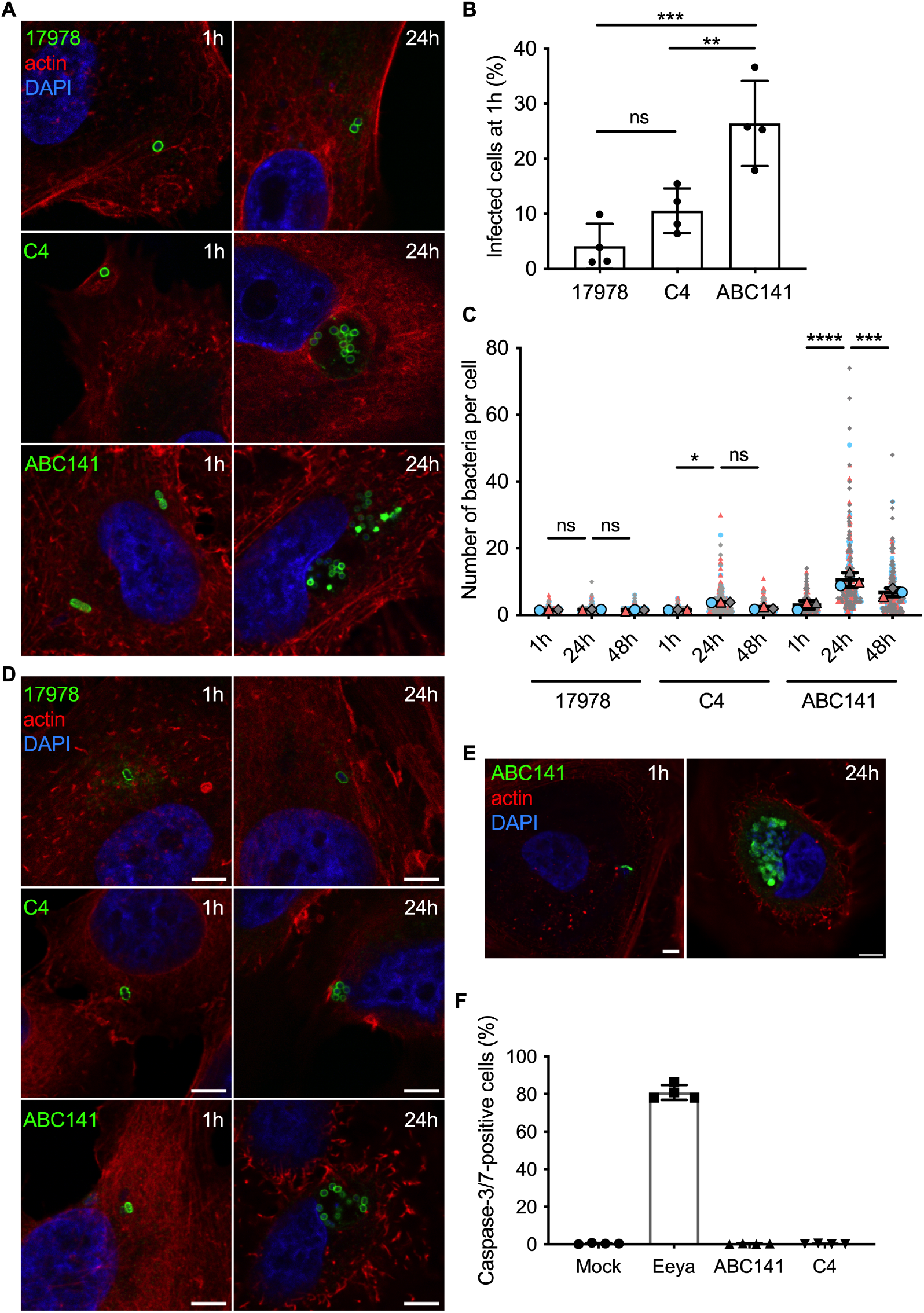
*A. baumannii* C4 and ABC141 strains multiply intracellularly. Human cells were infected with *A. baumannii* strains ATCC 17978, C4 or ABC141 and analyzed using confocal immunofluorescence microscopy. **(A)** Infected A549 cells were fixed at 1 h and 24 h p.i. and labelled with phalloidin and DAPI to visualize actin cytoskeleton (red) and nucleus (blue), respectively. *A. baumannii* strains were labelled with specific antibodies (green). Representative images are shown. Scale bars correspond to 5 μm. **(B)** Percentage of A549 cells infected by *A. baumannii* strains ATCC 17978, C4 or ABC141. Data correspond to mean *±* SD, from 4 independent experiments, counted by confocal microscopy. One-way ANOVA with a Holm-Sidak’s correction was used for multiple comparisons. Between C4 and ABC141 (**) *p*<0.01, between *A. baumannii* strains ATCC 17978 and ABC141 (***) *p*<0.001 and between 17978 and C4 (ns) *p*>0.05. **(C)** The numbers of intracellular bacteria per cell were counted at 1, 24 and 48 h p.i. and represented in “Superplot”. Colors (gray, blue and pink) correspond to 3 independent experiments. Each cell counted is shown (small dots) together with the means of each experiment (larger dots), which were used for statistical analysis. Statistical comparison was done with Two-way ANOVA with a Holm-Sidak’s correction for multiple comparisons. Between *A. baumannii* strains ATCC 17978 1 h and 24 h and ATCC 17978 24 h and 48 h (ns) *p*>0.05 for both comparisons, between C4 1 h and 24 h (*) *p*<0.05, between C4 24 h and 48 h (ns) *p*>0.05, between ABC141 1 h and 24 h (****) *p*<0.0001 and between ABC141 24 h and 48 h (***) *p*<0.001. **(D)** Infected EA.hy 926 cells were fixed at 1 and 24 h p.i. and labeled with phalloidin and DAPI to visualize actin cytoskeleton (red) and nucleus (blue), respectively. *A. baumannii* strains were labelled with specific antibodies (green). Representative images are shown. Scale bars correspond to 5 μm. **(E)** Human primary keratinocytes were infected with ABC141 for 24 h and immunolabelled with phalloidin (red), DAPI (blue) and an anti-*Acinetobacter* antibody (green). Scale bars correspond to 5 μm. **(F)** Quantification of the percentage of A549 cells that are caspase3/7-positive cells following infection with C4 or ABC141 for 24 h. Non infected cells were included as a negative control and cells treated incubated with Eeyarestatin for 24 h were used as positive control. C4 and ABC141 do not induce caspase-dependent cell death. Data correspond to mean *±* SD, from 4 independent experiments.

We next expanded our study to another clinical isolate available in the laboratory, the *A. baumannii* ABC141 strain, that represents IC5 and was isolated from a skin swab. A much higher percentage of cells were infected with *A. baumannii* ABC141 at 1 h p.i compared to C4 and ATCC 17978 (Fig. 2B), suggesting ABC141 is hyperinvasive. Also, at 24 h p.i, we observed a significant increase in the number of bacteria per cell (Fig. 2A and C) indicative of extensive intracellular replication. Interestingly, we often visualized multiple bacterial clusters per cell (Fig. 2A), with some clusters containing up to 50 bacteria. At 48 h p.i., however, a decrease in the numbers of ABC141 clusters was observed (Fig. 2C). It is important to note that when A549 cells were infected with a stationary phase culture of ABC141, we did not observe hyperinvasion nor significant intracellular replication suggesting that the growth stage is critical to confer hyperinvasive and replicative phenotypes to this strain, in contrast to the *A. baumannii* C4 strain.

To determine whether *A. baumannii* strains C4 and ABC141 are able to multiply within other cell types, we infected human endothelial EA.hy 926 cells. Similarly to A549 cells, all three strains invaded EA.hy 926 cells, with a few bacteria per cell visible at 1 h p.i. but only *A. baumannii* C4 and ABC141 were able to multiply intracellularly (Fig. 2D). We next infected primary human keratinocytes with the most invasive *A. baumannii* strain ABC141, to test its ability to multiply intracellularly in primary cells rather than immortalized cells lines. Extensive ABC141 multiplication was observed, confirming this phenotype is not cell type specific (Fig. 2E).

Because of the high numbers of intracellular bacteria observed for *A. baumannii* ABC141 at 24 h p.i., we next investigated whether heavily infected cells displayed signs of cell death. As previous reports suggest that *A. baumannii* can induce apoptosis (22–24), we monitored caspase 3 and 7 activation by microscopy at 24 h p.i. in comparison to mock-infected cells, and cells treated with high concentration of eeyarestatin as a positive control. Neither *A. baumannii* C4 nor ABC141 induced caspase-dependent cell death in A549 cells (Fig. 2F).

### Intracellular replicative strains of *A. baumannii* multiply within large non-acidic vacuoles positive for LAMP1

We next set out to characterize the nature of these *A. baumannii* intracellular compartments. We first labelled infected cells with the non-specific lectin Wheat germ agglutinin FITC conjugate (WGA-FITC) to visualize cellular membranes. This fluorescent probe labels sialic acid and glycoproteins containing β(1→4)-N-acetyl-D-glucosamine, such as cellulose, chitin and peptidoglycans. The majority of the intracellular bacterial clusters of *A. baumannii* C4 and ABC141 were surrounded by a WGA-positive membrane in A549 epithelial cells (Fig. 3A), suggesting enrichment in host cell sialic acid and/or bacterial glycoproteins on the vacuolar membrane surrounding replicating bacteria. To further characterize these *Acinetobacter* containing vacuoles (ACVs), we labelled A549 infected cells for the late endosomal/lysosomal marker LAMP1 (Lysosome-associated membrane glycoprotein 1). For both *A. baumannii* C4 and ABC141, ACVs were decorated with LAMP1 suggesting these are late endosomal or lysosomal-derived vacuoles (Fig. 3B). The proportion of LAMP-positive ACVs was quantified using the most invasive strain ABC141. Over 75% of ACVs were positive for WGA and LAMP1, confirming the vast majority of intracellular multiplying bacteria are within membrane-bound compartments (Fig. 3C). However, we did not detect any lysotracker associated with these ACVs, suggesting that this strain is inhibiting acidification or full lysosomal fusion (Fig. 3D). Of note, 13% of intracellular bacteria clusters were not labelled (Fig. 3C), which could correspond to cytosolic bacteria.

**Fig. 3.**
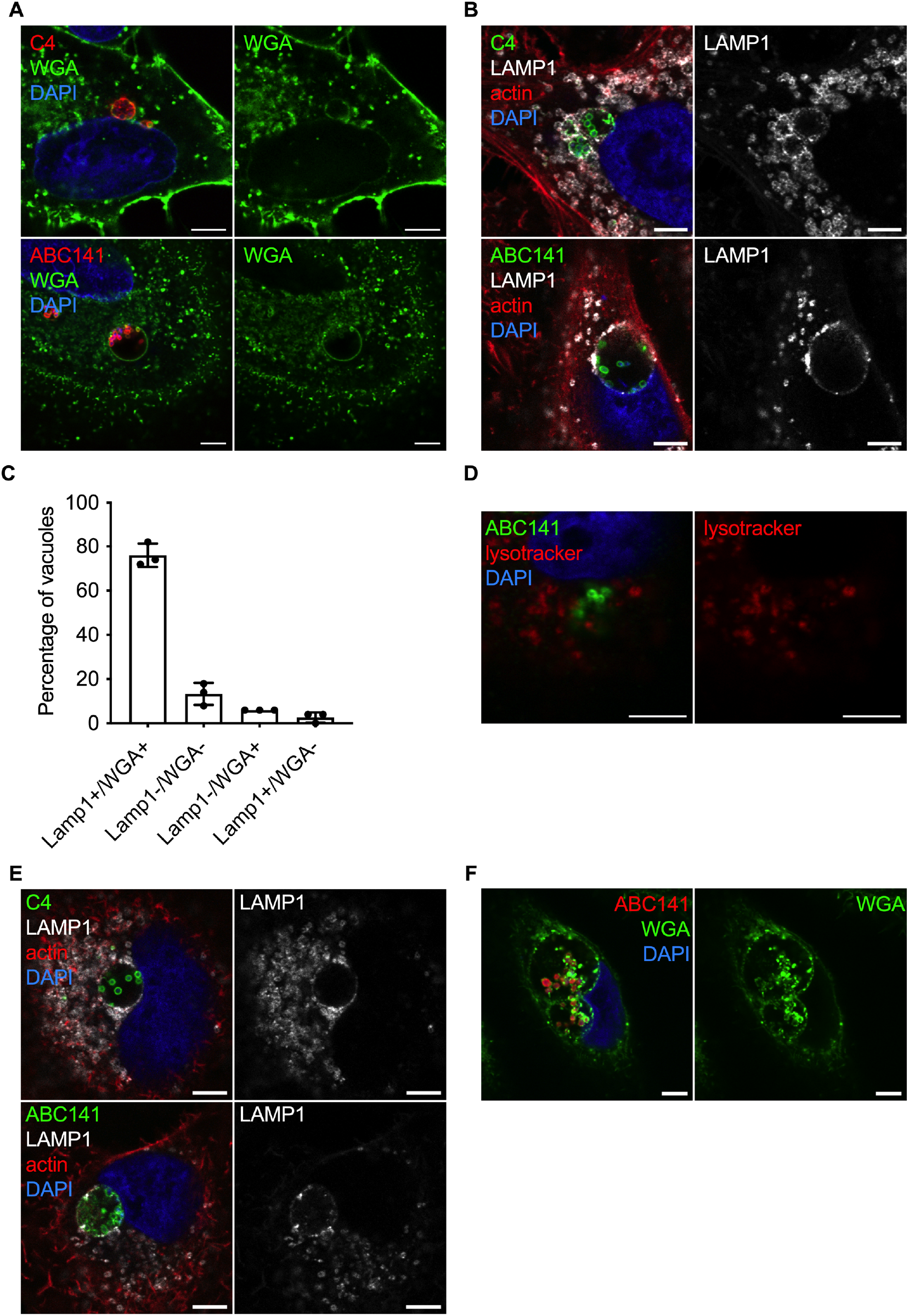
The multiplication of *A. baumannii* C4 and ABC141 occurs in large vacuoles positive for LAMP1. Human cells were infected with *A. baumannii* C4 or ABC141, immunolabelled 24h p.i. and analyzed using confocal immunofluorescence microscopy. Representative images are shown. Scale bars correspond to 5 μm. **(A)** Membranes of A549 cells were labelled with WGA (Green), the nucleus with DAPI (blue) and *A. baumannii* with antibodies (red). C4 and ABC141 multiply in *Acinetobacter*-containing vacuoles (ACV). **(B)** A549 cells were labelled with anti-LAMP1 antibodies (grey), with phalloidin and DAPI to visualize actin cytoskeleton and nucleus, respectively, and *A. baumannii* isolates were labelled with specific antibodies (green). ACV are positive for LAMP1 in A549 cells. **(C)** The number of ACV positive for LAMP-1 and/or WGA staining were counted at 24 h p.i. for cells infected by ABC141. ACV positive for both LAMP1 and WGA staining represent 78% of intracellular bacteria clusters. Data correspond to mean *±* SD, from 3 independent experiments. **(D)** A549 cells infected by ABC141 were labelled with Lysotracker DND-99 (red) 24h post-infection to visualize acidic compartments. ACVs are not acidic. **(E)** EA.hy 926 cells were labelled with anti-LAMP1 antibody (grey), anti-*A. baumannii* specific antibodies (green) and with phalloidin and DAPI to visualize actin cytoskeleton and nucleus. ACV are positive to LAMP1 in EA.hy 926 cells. **(F)** Representative images of human primary keratinocytes infected with ABC141 for 24 h. Nuclei were labelled with DAPI (blue), ABC141 were labelled with a specific antibody (red) and membranes with WGA (green).

To assess if *A. baumannii* C4 and ABC141 ACVs were WGA- and LAMP1-positive in other cell types, we infected EA.hy 926 cells and primary keratinocytes for 24 h (Fig. 3E and F). Microscopic analysis confirmed that both strains multiply in equivalent intracellular compartments in endothelial cells and primary keratinocytes. Indeed, *A. baumannii* C4 and ABC141 multiply in very large vacuoles whose size could exceed 7 μm, in some cases seemingly “pressing” against the nucleus (Fig. 3E and F). Taken together, these results indicate that the intracellular replicative strains of *A. baumannii* C4 and ABC141 multiply in LAMP1-positive non-acidic vacuoles in human non-phagocytic cells.

### *Acinetobacter*-containing vacuoles have a single membrane and do not colocalize with the autophagy marker LC3

In view of the large size of ACVs and the presence of LAMP1 we next hypothesized that these could correspond to autophagosomes, as previously demonstrated surrounding intracellular *A. baumannii* strain AB5075 before bacterial killing (17). To test this hypothesis, we immunolabelled A549 infected cells for the microtubule-associated protein 1A/1B-light chain 3 (LC3), widely used as a marker of autophagy. We did not observe any LC3 labelling associated with *A. baumannii* C4 or ABC141 ACVs (Fig. 4A). To confirm that ACVs were not autophagy-derived vacuoles, we performed transmission electron microscopy on A549 cells infected with ABC141, which shows a higher rate of invasion. We observed *A. baumannii* ABC141 exclusively within large vacuoles composed of a single membrane, confirming that ACVs are not autophagosomes. Moreover, we noted that ACVs contain free space and multiple small vesicles (Fig. 4B). In summary, we found that intracellular replicative strains of *A. baumannii* can create a niche favorable to their multiplication within human cells.

**Fig. 4.**
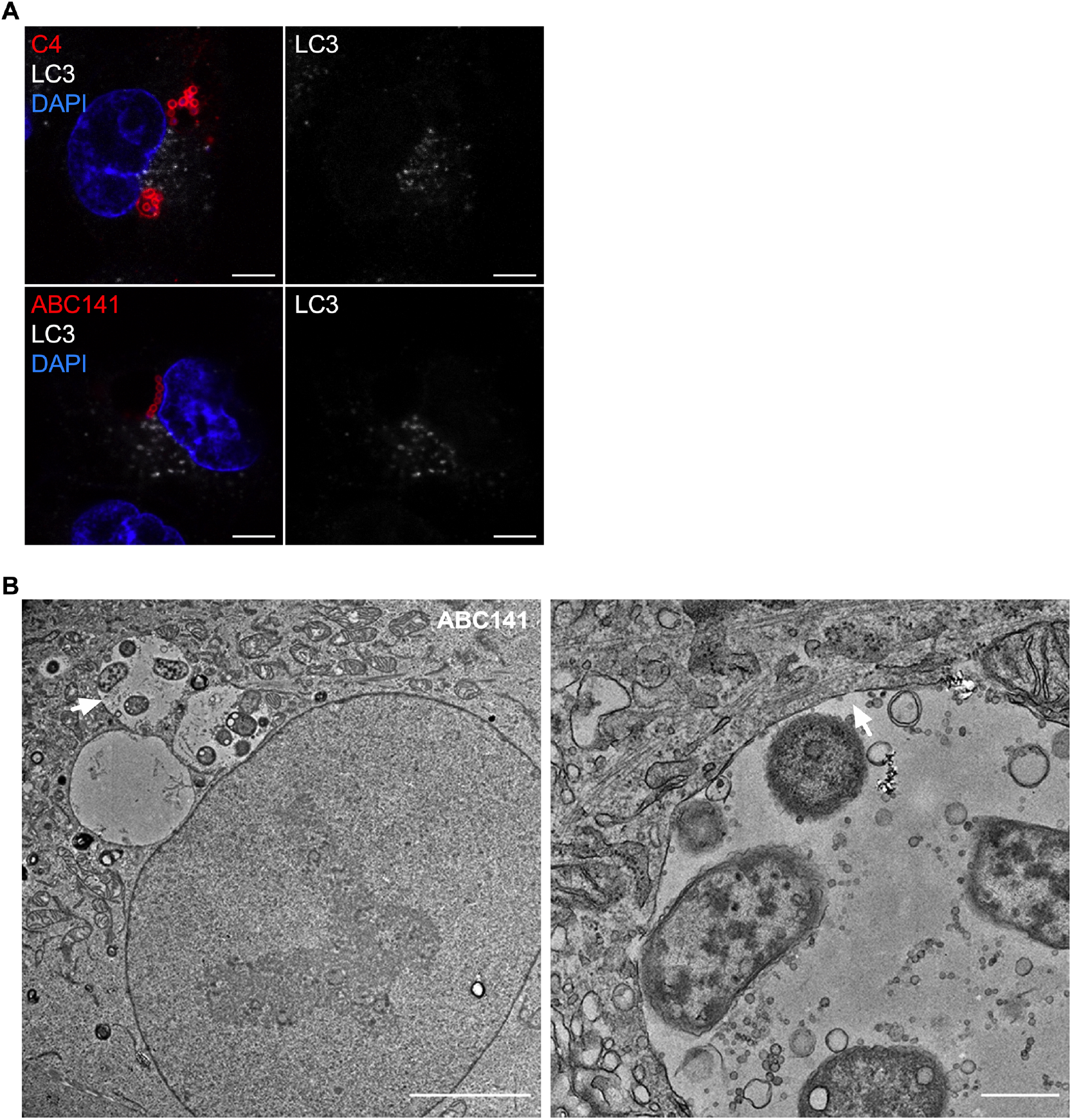
ACV are single-membrane vacuoles that do not colocalize with the autophagy marker LC3. **(A)**A549 cells were infected with *A. baumannii* C4 or ABC141, immunolabelled 24h post-infection and analyzed using confocal immunofluorescence microscopy. *A. baumannii* (red) and LC3 (white) were labelled with antibodies and nucleus with DAPI (blue). ACV do not colocalize with LC3. Representative images are shown. Scale bars correspond to 5 μm. **(B)** Transmission electron microscopy of infected A549 cells by ABC141 24 h post-infection. The scale bar corresponds to 5 μm. The second picture represents a zoom of the vacuole indicated by the white arrow. The scale bar corresponds to 500 nm. ABC141 multiplies in ACV with a single membrane.

### Prevalence of intracellular replicative strains in clinical *A. baumannii* isolates

Since with this small selection of *A. baumannii* clinical isolates we observed several distinct phenotypes, we gathered a larger collection of 43 non-duplicate clinical isolates to determine the prevalence of each phenotype. IC4 and IC5 isolates were chosen to compare with C4 and ABC141, respectively, while other clonal lineages and non-lineages were chosen to compare across the lineages (Figure 5, Supplementary Table 1). To screen a high number of isolates we set up a high-content screening, to image infected A549 cells at 24 h p.i. labelled for LAMP1 to clearly distinguish intracellular bacterial clusters in combination with a mix of three antibodies against *A. baumannii*, which we confirmed beforehand could label all bacteria tested. We classified the observed phenotypes in 4 categories: 1) non-invasive; 2) capable of entry and survival; 3) capable of replication, visible by the formation of LAMP1-positive bacterial clusters and 4) hyperinvasive, with a rate of infection equivalent to the *A. baumannii* ABC141 strain. In total, 46 isolates were screened in two independent assays, including ATCC17978, C4 and ABC141. From this collection, 24 isolates were non-invasive, with no bacteria detected intracellularly, whereas 18 isolates were capable of cell entry but did not show any intracellular multiplication, giving a phenotype equivalent to the *A. baumannii* ATCC 17978. The four strains C4, ABC141, BMBF193 and R10 were capable of intracellular multiplication in LAMP1-positive vacuoles but only the ABC141 and BMB193 were also hyperinvasive. These results highlight the variety of phenotypes observed for clinical *A. baumannii* strains and confirm that a subset of these have the ability to invade and multiply within host cells.

**Fig. 5.**
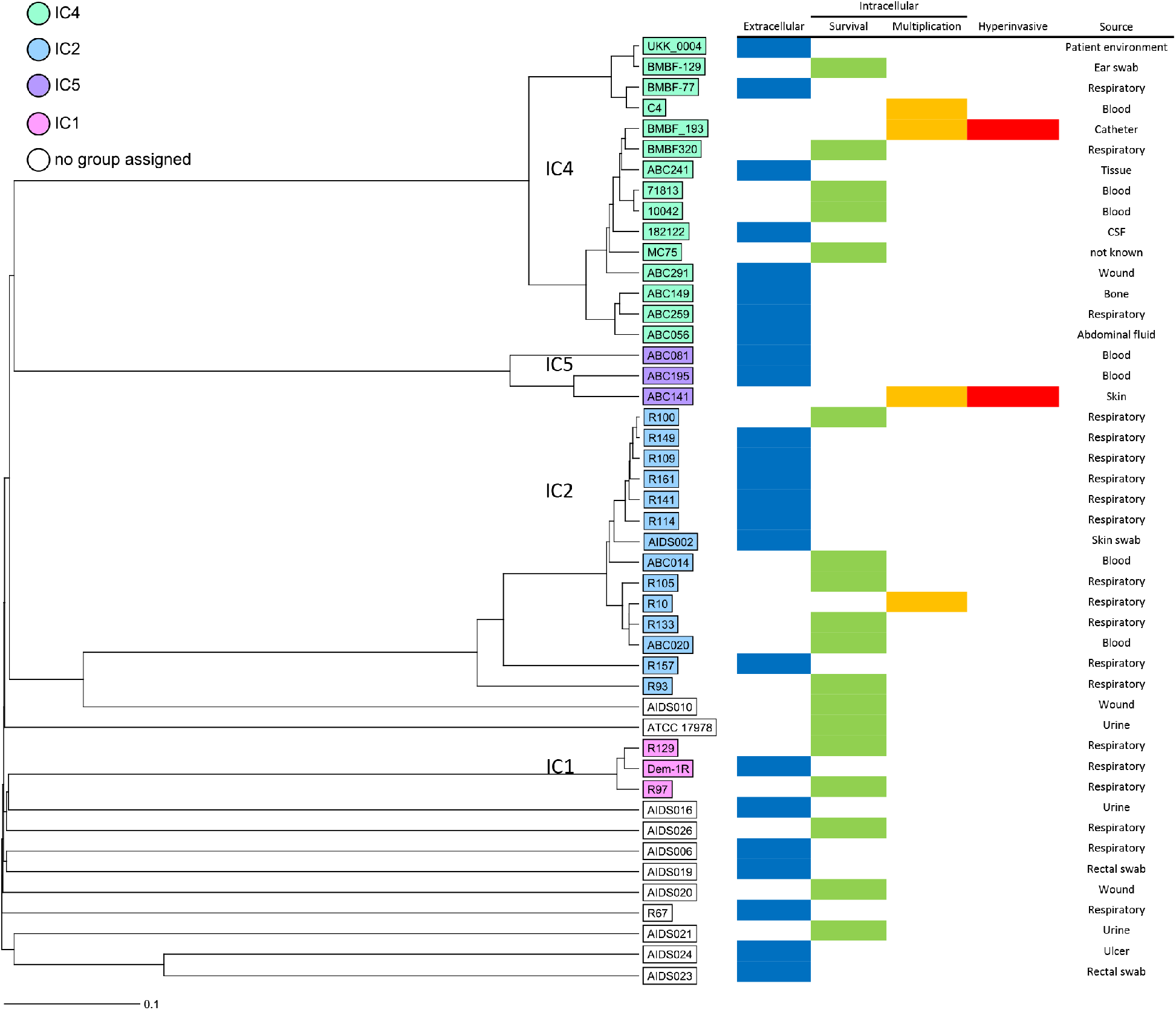
UPGMA tree generated using cgMLST data showing the results of a high-content screen by confocal microscopy of clinical *A. baumannii* isolates. A549 cells were infected with 43 clinical *A. baumannii* isolates and *A. baumannii* C4, ABC141 and ATCC 17978 strains as controls. Cells were fixed at 24 h p.i. and immunolabelled with a mix of three anti-*Acinetobacter* antibodies, LAMP1 and DAPI. For each strain, z-stacks series were imaged at five different positions of the well. cgMLST was performed using Ridom SeqSphere+ using 2390 targets. Isolates are coloured based on clustering with a clonal lineage. Those within white boxes do not cluster with any of the lineages. The source of the strains is indicated.

## Discussion

In this study we highlight a diversity of phenotypes exhibited by *A. baumannii* clinical isolates regarding their interaction with human non-phagocytic cells. The observed interactions were strain specific and there was no correlation within lineages.

A subset of these strains, including one that was associated with high fatality rate, are capable of active intracellular replication without induction of cytotoxicity. This highlights an important niche that could be hindering treatment of patients infected with these types of strains.

Although considered originally as an extracellular pathogen, a growing number studies have shown that some strains of *A. baumannii* are able to invade epithelial and endothelial human cells (13–20, 23, 25–28). *A. baumannii* cell invasion occurs via a zipper-like mechanism involving actin micro-filaments and microtubules (13, 19), and is dependent on clathrin, β arrestins and phospholipase C-coupled G-proteins, as their inhibition prevents its entry into the cell (26). The platelet activating factor receptor (PAFR) was also shown to enhance bacterial internalization following interaction with the phosphorylcholine-containing outer membrane protein of *A. baumannii* (26). However, this interaction results in an increase in intracellular calcium concentration, and subsequent cell death. The use of random mutant libraries highlighted the importance of *A. baumannii* phospholipases in cellular invasion (25). The outer membrane proteins OmpA and Omp33 of *A. baumannii* have also been implicated but the cellular interactions taking place remain unknown (13, 19). A diversity of phenotypes regarding the ability to invade cells has been previously reported (16). The discovery of hyperinvasive strains of *A. baumannii* described in our study will provide an excellent tool to decipher the underlying mechanisms. All of these bacterial factors implicated in internalization are encoded within the genome of *A. baumannii* ABC141, suggesting enhanced invasion could be due to different regulatory mechanisms, or alternatively, to the presence of new bacterial factors involved yet to be identified. Detailed genome and transcriptome comparisons are necessary to decipher the virulence factors involved.

Although all these data strongly support the existence of an *A. baumannii* intracellular phase, the fate of intracellular bacteria remains less well characterized. Intracellular persistence of *A. baumannii* in cultured cells was described in both epithelial cells and macrophages (11, 13–17, 18). In most cases, bacteria were found to transiently survive within an acidic and autophagy-derived compartment, which eventually results in efficient bacterial killing (16). In our study we found several *A. baumannii* clinical strains, capable of extensive intracellular multiplication for up to 24 h and 48 h p.i. Importantly, these observations were done by microscopy which allowed us to ensure the increase in bacterial numbers is not due to extracellular replication of bacteria that are not efficiently killed by the antibiotics. We have observed for a few antibiotic sensitive and resistant strains excluded from this study, the formation of bacterial aggregates tightly adhered to the surface of the cells and the coverslips, that seem protected from antibiotic treatment, which would mislead CFU counts. Interestingly, other studies have identified clinical strains able to multiply in cultured macrophages by bacterial CFU counts (21), suggesting these phenotypes could be extended to phagocytic cells. Consistently, while this work was being submitted, a study has shown that *A. baumannii* urinary tract infection strains were capable of intracellular replication in cultured macrophages (29), by both CFU and microscopy counts, strengthening the hypothesis that an intracellular replication niche may be clinically relevant for a subset of *A. baumannii* strains.

In this study we found that the *A. baumannii* C4 and ABC141 multiply within membrane-bound vacuoles, reaching 20 to 70 bacteria at 24 h p.i., not previously observed in *A. baumannii* infected cells. ACVs are very large compartments, surrounded by a glycoprotein rich membrane which colocalizes with LAMP1. Further work has to be done to determine if the glycoproteins decorating ACVs are from eukaryotic or bacterial origin, or both. The presence of LAMP1, indicative of a lysosomal-derived compartment was not accompanied by acidification as we did not detect any Lysotracker positive ACVs. These results are in contrast with previous reports showing persistence of some *A. baumannii* within LAMP1 acidic compartments, notably for the *A. baumannii* 19606 strain (20). It has been shown that *A. baumannii* multiply slower in acidic pH environment *in vitro* and the inhibition of acidification by bafilomycin A1 increase the number of intracellular *A. baumannii* AB5075 (17, 18). Therefore, we hypothesize that our *A. baumannii* clinical strains are able to establish a niche suitable for intracellular replication by preventing fusion with degradative lysosomes and blocking their acidification. Interestingly, these vacuoles attain very large sizes even pushing the nucleus of the cell. To our knowledge, this type of vacuole is not commonly observed for intracellular pathogens and merits further investigation. One example of a pathogen reported to create large multivesicular vacuoles with the appearance of “empty space” reminiscent of these ACVs is *Helicobacter pylori* (30). This pathogen colonizes human stomach mucosa, causes gastrointestinal diseases and multiplies in human cells. It has been shown that intracellular *H. pylori* can modulate autophagy, blocking the acidification of the vacuoles and multiplying inside large autophagosomes (31).

Autophagy is a key cellular process for cell survival and host innate immunity that participates in the elimination of invading bacteria. Not surprisingly, many pathogens have developed mechanisms to modulate autophagy or hijack this process in order to promote their survival and multiplication. In the case of *A. baumannii* infection, there are a few reports implicating autophagy during infection. Indeed, the *A. baumannii* ATCC 17978 strain was reported to persist within double-membrane vacuoles (23). The porin Omp33-36 was the virulence factor implicated in induction of apoptosis via caspase activation and induction of autophagy visible by the accumulation of p62 and LC3B-II (23). *A. baumannii* infection itself was shown to induce Beclin-1 dependent autophagy via the AMPK/ERK signaling pathway (20), with the involvement the porin OmpA (15). In addition, the transcriptional factor EB (18) was shown to block acidification of the autophagosome-lysosome system and enhance bacterial persistence. In contrast to all these reports, ACVs described in this study did not show any LC3 immunolabelling. Furthermore, transmission electron microscopy confirmed that ACVs are single-membrane vacuoles and therefore not autophagic in nature. We conclude that the *A. baumannii* clinical strains capable of intracellular multiplication presented in this study create specialized replicative vacuoles that successfully escape autophagy and subsequent lysosomal degradation. After 24h of replication, in the case of the *A. baumannii* ABC141 strain, we observed a reduction in the presence of large vacuoles. The recent work of the Feldman lab showed that *A. baumannii* urinary tract infection strains capable of intra-macrophage multiplication (29), escape from the cell. So, it is likely that a similar event is taking place for *A. baumannii* C4 and ABC141 in epithelial cells, allowing bacteria to multiply without killing the host cell and then egress from infected cells to disseminate within the tissue or systemically.

The intracellular replication phenotype we observed seems to be common to several clinical isolates, suggesting this phenotype is relevant from a clinical point of view. In addition to the 4 initial strains studies, 43 strains were tested using a high-content microscopy screen, allowing us to get some insight into the prevalence of this phenotype. We found that the ability to invade and persist without multiplication was quite frequent amongst the different isolates tested, but this was irrespective of their lineages, suggesting strain specific traits that have yet to be identified. Importantly, 4 strains in total in our study were capable of intracellular replication, all within large LAMP1-positive vacuoles, suggesting that a significant proportion of current *A. baumannii* isolates share this intracellular niche. It is important to keep in mind that we may be under-estimating the prevalence of this phenotype due to the specificity of the conditions tested. For example, some *A. baumannii* may have tissue specificity or require particular growth conditions. Consistently, an overnight culture of *A. baumannii* ABC141 significantly reduces its invasion capacity while this is not the case for *A. baumannii* C4.

Interestingly, intracellular multiplication was observed for both the virulent *A. baumannii* C4 isolate, but also for *A. baumannii* ABC141 which was not isolated from an infection site. This result suggests that the ability to establish an intracellular niche is not the direct cause of enhanced virulence in patients. This is not surprising as the gravity of an *A. baumannii* infection is tightly connected to the host susceptibility and hence it is not possible to assign levels of virulence to different *A. baumannii* strains based on the site of isolation. It is possible that virulence properties are randomly distributed among clinical isolates and their presence or absence does not translate into clinical pathogenicity. Nonetheless, an intracellular phase could confer enhanced protection to the pathogen against some antibiotics, could promote the dissemination in the organism or give rise to relapsing infections. This phenomenon has been described in another nosocomial pathogen, *Staphylococcus aureus*, which is also able to survive and multiply inside host cells (32, 33). The invasion of human THP-1 macrophages protects *S. aureus* from vancomycin, oxacillin, moxifloxacin, rifampicin, gentamicin and oritavancin (34). In a mouse model, intracellular *S. aureus* can establish an infection even in the presence of vancomycin (35). Furthermore, *S. aureus* internalized in keratinocytes are not killed by antibiotics even at 20 fold their minimal inhibitory concentration (36). Similar observations were made with invasive *P. aeruginosa* strains, that are protected from aminoglycosides in contrast to ß-lactams, fluoroquinolones, or colistin (37). Therefore, the intracellular nature of specific *A. baumannii* strains may have important clinical consequences. The protection conferred by the intracellular environment could aggravate antibiotic resistance of *A. baumannii* in patients which will not be detected by antibiotic susceptibility testing *in vitro*. Discriminating intracellular replicating strains may provide an important diagnostic tool in the future. Our work is a first step in the identification of hyperinvasive strains and replicative strains, providing insight on their intracellular trafficking which may ultimately prove beneficial to help adapt antimicrobial therapies against this nosocomial pathogen.

## Materials and Methods

### Culture of cell lines and primary cells

A549 (human epithelial lung cell line) and EA.hy926 (human endothelial somatic cell line) cells were bought from Merck company and ATCC, respectively. Both cell lines were grown in Dulbecco’s modified Eagle medium (DMEM) supplemented with 1% L-glutamine and 10% of fetal calf serum at 37°C with 5% CO2 atmosphere. Normal human keratinocytes (NHK) cultures were established from foreskin after dermal-epidermal dissociation with 0.05% trypsin and 0.01% ethylenediaminetetraacetic acid (EDTA) in phosphate-buffered saline (PBS), pH 7.4 and grown in supplemented keratinocyte growth medium containing 0.15 mM CaCl2 (KBM-2 BulletKit, Lonza Biosciences, Basel, Switzerland) as previously described (38). NHK were used between passages 1-3. Biopsies were obtained following ethical and safety guidelines according to French regulation donors (Declaration no. DC-2008-162 delivered to the Cell and Tissue Bank of Hospices Civils de Lyon).

### Bacterial strains and culture conditions

Bacterial strains, culture conditions and whole genome sequencing The ATCC 17978 strain was obtained from ATCC and all other clinical isolates of *A. baumannii* were provided by Harald Seifert and Paul Higgins from the University of Cologne, Germany (Supplemental Table 1). *A. baumannii* strains were grown on LB agar pH 7.4 for 24 h at 37°C. For liquid cultures, a single colony was inoculated in LB pH 7.4 and incubated for 17 h for stationary phase cultures, for the experiments described in Figure 1. All other experiments were done with late exponential cultures, in which a colony was inoculated in LB overnight (16 h) and then diluted 1/100 and grown until an OD600 of 0.8-1. Whole genome sequencing and analysis was performed as previously described (39). The raw sequencing reads used in this project were submitted to the European Nucleotide Archive (https://www.ebi.ac.uk/ena/) and their nucleotide accession numbers are listed in Supplementary Table 1.

### Human cells infection and adhesion assays

Human cells were grown in 96-well culture plates at 1.6×10^4^ cells/well, 24-well culture plates at 4 × 10^4^ cells/well, 12-well culture plates at 2,5 × 10^5^ cells/well and 6-well-culture plates at 1 × 10^6^ cells/well for the high throughput screen, fluorescent microscopy, adhesion assays and electron microscopy, respectively. Then, they were infected at a multiplicity of infection (MOI) of 100 of *A. baumannii* in supplemented DMEM medium pre-warmed at 37 °C. Plates were centrifuged at 400g for 5 min and incubated 1 h at 37 °C with 5% CO2 atmosphere. Once incubated, plates were washed 5 times with PBS to remove extracellular bacteria. For adhesion assays: Cells were lysed by incubation for 5 min with 0.1% sodium deoxycholate. Bacteria were enumerated before and after infection to calculate the percentage of adhesion. The MOI was verified by the number of CFU/ml in the inocula. For fluorescent microscopy: Infected cells were incubated in supplemented DMEM medium with apramycin (40 μg/ml) or tobramycin (50 μg/ml) for 24 h or 48 h. After 24h of incubation, the antibiotic was changed followed by a second incubation of 24 h at 37 °C with 5% CO2 atmosphere (time-point 48h). Coverslips were fixed at each time point (1 h, 24 h and 48 h) with Antigenfix (Diapath, paraformaldehyde pH 7.2-7.4) for 15 min or methanol (pre-cooled at −20 °C) for 5 min at room temperature. Finally, samples were washed with PBS 5 times and kept at 4 °C. For electron microscopy: Samples were fixed in 2% glutaraldehyde and 2% paraformaldehyde in 0.1 M cacodylate buffer (pH 7.2) overnight at 4 °C. After extensive washing in 0.1 M cacodylate at 4 °C, cells were post-fixed with 2% osmium tetroxide in 0.1 M cacodylate buffer for 1 h at 4 °C. After rinsing with water, a contrast was performed with 1% uranyl acetate. The samples were then dehydrated using a graded series of ethanol and embedded in Epon resin. After polymerization at 60 °C for 48 h, ultrathin sections (60 nm) were cut using a Leica UC7 microtome and contrasted with lead citrate. Samples were examined with a JEOL 1400 Flash transmission electron microscope.

### Immunolabelling

Once fixed, cells were incubated in PBS with 1% saponin and 2% Bovine Serum Albumin (BSA) for 1 h at room temperature for permeabilization and blocking. Primary antibodies were diluted in the blocking solution and incubated for 2 h. To label bacteria we raised antibodies in rabbits against *A. baumannii* ATCC 17978, AB5075 and C4, using a lyophilized preparation and a speedy-immunization polyclonal program (BIOTEM, France; Eurogentec, Belgium). The anti-*A. baumannii* antibody mix was diluted at 1/1000 for clinical strains and at 1/10,000 for the ATCC 17978 strain. The following additional antibodies were used: mouse anti-LC3 (1/1000), mouse anti-βtubulin (1/200), mouse anti-LAMP1 H4A3 (1/200), from the Developmental Studies Hybridoma Bank, created by the NICHD of the NIH and maintained at the University of Iowa. Coverslips were then washed twice in PBS, 0.1% saponin, BSA 2%. Secondary antibodies and dyes, anti-rabbit Alexa-488 or 555 (1/500), anti-mouse Alexa-555 or 647 (1/500), Wheat Germ Agglutinin (WGA) FITC conjugate (Sigma, 1/200), Phalloïdine 647N ATTO (1/200) and DAPI nuclear dye (Bio-rad,1/1000) were diluted in the blocking solution. Cells were incubated for 1 h followed by two washes in PBS with 0.1% saponin/BSA 2%, one wash in PBS and one wash in distilled water. Finally, coverslips were mounted with ProLong Gold (ThermoFisher). For the high-content screening cells were labeled with LAMP1 and a mix of the 3 home-made antibodies used in this study, which enabled labeling of all the strains. DAPI was also included.

### Caspase 3/7 detection

A549 cells were infected for 24 h as described above. Non-infected cells incubated with Eeyarestatine (500 μmM) for 24h were used as positive control. Each condition was next incubated with the Caspase 3/7 Green detection reagent (CellEventTM Kit) diluted in PBS with 5% FCS, for 30 minutes. Cells were fixed with Antigenfix for 15 min at room temperature. Finally, samples were washed with PBS 5 times and kept at 4°C.

### Lysotracker

A549 cells were infected for 22 h as described above. They were then incubated for 2 h with the Lysotracker DND-99 (75 nM) in DMEM medium pre-warmed at 37°C. Finally, the loading solution was replaced by Antigenfix to fix cells. Labelled cells were immediately observed by confocal microscopy.

### Counting the number of infected cells and intracellular bacteria

The percentage of infected cells at 1 or 24 h post-infection was calculated by counting the number of non-infected and infected cells per 10 fields for *A. baumannii* ATCC 17978, C4, ABC141 and ABC56 for 3 independent experiments. The number of intracellular bacteria per cell for 17978, C4 and ABC141 strains were counted for 3 independent experiments. Results are represented in “superplot” as described by Lord and colleagues (40), in which each experiment is color-coded. All the cells counted are presented as the corresponding mean and standard deviation. Statistical analysis was done by comparing the means of independent experiments. The number of vacuoles with a LAMP1 and/or WGA labelling were counted for *A. baumannii* ABC141 in 3 independent experiments.

### Confocal microscopy

For all images and counting, coverslips were mounted with ProLong Gold (ThermoFisher) and observed with a Zeiss LSM800 Airy Scan laser scanning confocal microscope with an oil immersion objective 63x. For the high throughput screen, images were collected with a Yokogawa CQ1 and a 40x objective was used. Finally, they were analyzed with FIJI (41) and assembled in FigureJ (42).

### Experimental infection (*Galleria*)

G. mellonella larvae, purchased from Sud Est Appats (http://www.sudestappats.fr/), were used within 48 h of arrival. *A. baumannii* isolates tested were grown overnight in LB and then diluted with PBS to enable injection of 1×10^6^ CFU, as determined by CFU plating. Bacterial suspensions were injected into the hemolymph of each larva (second last left proleg) using a Hamilton syringe (10 μl). Groups of twenty randomly picked larvae were used for each strain. Survival curves were plotted using Graphpad, and comparisons in survival were calculated using the log-rank Mantel-Cox test.

### IL-6 quantifications

A549 cells were grown in 96-well culture plates at a density of 1×10^5^ cells/mL. The next day, the cells were infected with the different *A. baumannii* isolates from a culture in stationary phase diluted to an MOI of 100. The infection protocol is the same as described above. The supernatants were recovered after 8, 24 or 48 h of infection. The concentration of IL-6 was quantified by ELISA (Human IL-6 ELISA Ready-SET-Go!, Thermofisher) by following the supplier’s protocol.

### Statistical tests

All data sets were tested for normality using Shapiro-Wilkinson test. When a normal distribution was confirmed we used a One-Way ANOVA test with a Holm-Sidak’s correction for multiple comparisons. For two independent variables, a Two-Way ANOVA test was used. For data sets that did not show normality, a Kruskall-Wallis test was applied, with Dunn’s correction. All analyses were done using Prism Graph Pad 7.

## ACKNOWLEDGEMENTS

We acknowledge the contribution of the SFR Biosciences (UAR3444/CNRS, US8/Inserm, ENS de Lyon, UCBL) imaging facility Plateau Technique Imagerie/Microcopie (PLATIM) and the Centre Technologique des Microstructures, Université Lyon 1. We are grateful to Matthias Faure (CIRI, Lyon) for providing us with the anti-LC3 antibody and advice. This work was funded by the Fondation pour la Recherche Médicale grant DEQ20180339215. S. Salcedo is an INSERM researcher. S. Göttig was supported by the Deutsche Forschungsgemeinschaft (DFG FOR 2251).

## Author contributions

T. Rubio carried out and analyzed all experiments with *A. baumannii* ABC141, and quantifications of intracellular multiplication and trafficking of all isolates. S. Gagné characterized the C4 strain and discovered the intracellular multiplication phenotype. C. Debruyne carried out and analyzed the caspase experiments and high-content screening. D. Mongellaz set up and managed the clinical collection and optimized the experimental protocol for the screening. C. Dias assisted T. Rubio with infection experiments. C. Cluzel did the electron microscopy and P. Rousselle cultured and provided the keratinocytes. H. Seifert, P. Higgins and S. Göttig provided all the clinical isolates and input regarding the epidemiology, genomics and interpretation of the screen results. The project was conceived, supervised and funded by S. Salcedo. The manuscript was written by T. Rubio and S. Salcedo. All authors read an approved the manuscript. I would also like to thank R. Henriques (IGC, Portugal) for making the BioRxiv Overleaf Latex template used for formatting this manuscript.

---

**Supplementary Table 1.** Strains used in this study. IC; international clone, CSF; cerebrospinal fluid

| Strain | IC | Source | Nucleotide Accession | doi |
| --- | --- | --- | --- | --- |
| Dem1R | IC1 | Respiratory | ERR5709465 | doi:10.1128/AAC.00598-10 |
| R129 | IC1 | Respiratory | ERR2451960 | doi: 10.1093/jac/dkx322 |
| R97 | IC1 | Respiratory | ERR2455403 | doi: 10.1093/jac/dkx322 |
| ABC14 | IC2 | Blood | ERR2712988 | doi: 10.1093/jac/dkab079 |
| ABC20 | IC2 | Blood | ERR2711313 | doi: 10.1093/jac/dkab079 |
| AIDS-002 | IC2 | Skin swab | ERR5709471 | unpublished |
| R10 | IC2 | Respiratory | ERR2451540 | doi: 10.1093/jac/dkx322 |
| R100 | IC2 | Respiratory | ERR2451537 | doi: 10.1093/jac/dkx322 |
| R105 | IC2 | Respiratory | ERR2451538 | doi: 10.1093/jac/dkx322 |
| R109 | IC2 | Respiratory | ERR2451539 | doi: 10.1093/jac/dkx322 |
| R114 | IC2 | Respiratory | ERR2451921 | doi: 10.1093/jac/dkx322 |
| R133 | IC2 | Respiratory | ERR2451963 | doi: 10.1093/jac/dkx322 |
| R141 | IC2 | Respiratory | ERR2452000 | doi: 10.1093/jac/dkx322 |
| R149 | IC2 | Respiratory | ERR2452005 | doi: 10.1093/jac/dkx322 |
| R157 | IC2 | Respiratory | ERR2452101 | doi: 10.1093/jac/dkx322 |
| R161 | IC2 | Respiratory | ERR2455357 | doi: 10.1093/jac/dkx322 |
| R93 | IC2 | Respiratory | ERR2455413 | doi: 10.1093/jac/dkx322 |
| 10042 | IC4 | Blood | ERR5709468 | doi: 10.3389/fmicb.2020.584603 |
| 71813 | IC4 | Blood | ERR5709467 | doi: 10.3389/fmicb.2020.584603 |
| 182122 | IC4 | CSF | ERR5709469 | doi: 10.3389/fmicb.2020.584603 |
| ABC149 | IC4 | Bone | ERR2712285 | doi: 10.1093/jac/dkab079 |
| ABC241 | IC4 | Tissue | ERR2712376 | doi: 10.1093/jac/dkab079 |
| ABC259 | IC4 | Respiratory | ERR2712393 | doi: 10.1093/jac/dkab079 |
| ABC291 | IC4 | Wound | ERR4436640 | doi: 10.1093/jac/dkab079 |
| ABC56 | IC4 | Abdominal fluid | ERR2711689 | doi: 10.1093/jac/dkab079 |
| BMBF-129 | IC4 | Ear swab | ERR5709482 | doi:10.1093/jac/dkp428 |
| BMBF-193 | IC4 | Catheter | ERR5709481 | doi:10.1093/jac/dkp428 |
| BMBF-320 | IC4 | Respiratory | ERR5709466 | doi:10.1093/jac/dkp428 |
| BMBF-77 | IC4 | Respiratory | ERR5709502 | doi:10.1093/jac/dkp428 |
| C4 | IC4 | Blood | ERR5709472 | unpublished |
| MC75 | IC4 | not known | ERR5709464 | doi.org/10.1016/j.ijantimicag.2019.03.019 |
| UKK_0004 | IC4 | Patient environment | ERR1226902 | DOI 10.1099/jmm.0.012302-0 |
| ABC141 | IC5 | Skin | ERR2712278 | doi: 10.1093/jac/dkab079 |
| ABC195 | IC5 | Blood | ERR2712330 | doi: 10.1093/jac/dkab079 |
| ABC81 | IC5 | Blood | ERR2712148 | doi: 10.1093/jac/dkab079 |
| AIDS-021 | IC7 | Urine | ERR5709475 | unpublished |
| AIDS-006 | no | Respiratory | ERR5709470 | unpublished |
| AIDS-010 | no | Wound | ERR5709480 | unpublished |
| AIDS-016 | no | Urine | ERR5709476 | unpublished |
| AIDS-019 | no | Rectal swab | ERR5709474 | unpublished |
| AIDS-020 | no | Wound | ERR5709473 | unpublished |
| AIDS-023 | no | Rectal swab | ERR5709479 | unpublished |
| AIDS-024 | no | Ulcer | ERR5709477 | unpublished |
| AIDS-026 | no | Respiratory | ERR5709478 | unpublished |
| ATCC 17978 | no | Laboratory strain | NZ_CP012004.1 | unpublished |
| R67 | no | Respiratory | ERR2455376 | doi: 10.1093/jac/dkx322 |

